# Cost of resistance: an unreasonably expensive concept

**DOI:** 10.1101/276675

**Authors:** Thomas Lenormand, Noémie Harmand, Romain Gallet

## Abstract

This preprint has been reviewed and recommended by Peer Community In Evolutionary Biology (https://doi.org/10.24072/pci.evolbiol.100052). The cost of resistance, or the fitness effect of resistance mutation in absence of the drug, is a very widepsread concept in evolutionary genetics and beyond. It has represented an important addition to the simplistic view that resistance mutations should solely be considered as beneficial mutations. Yet, this concept also entails a series of serious difficulties in its definition, interpretation and current usage. In many cases, it may be simpler, clearer, and more insightful to study, measure and analyze the fitness effects of mutations across environments and to better distinguish those effects from ‘pleiotropic effects’ of those mutations.

## Introduction

The study of resistance to antibiotics, insecticides, acaricides, fungicides, herbicides, chemotherapy drugs etc. is, for obvious reasons, a very active field of research. We are in the middle of a “crisis,” which has important consequences for public health and agriculture (e.g. World Health Organization 2014; Ventola 2015). It is yet unclear whether we will be able to deal with “superbugs”, “superweeds” and other “supermicrobes” in the near future. Studies have focused intensely on the genetic, cellular, and biochemical mechanisms responsible for resistance, but also on the fitness effect of those mutations.

### Resistance mutations as beneficial mutations

Resistance evolution is a particular case of the more general situation of adaptation to new environmental conditions. Processes of adaptation have been intensely studied from Darwin’s time, since refined by powerful population genetic concepts that have been put forward in the modern synthesis (Orr 2005). In brief, resistance mutation are beneficial mutations in treated environments. They spread because they confer an obvious fitness advantage in presence of the drug (the antibiotic, insecticide, acaricide, fungicide etc.). By definition, a resistance mutation allows for survival while the susceptible wild-type simply dies when exposed to the drug. Hence, classically, the fitness benefit of a resistance mutation (relative to a susceptible one) depends on the fraction of the population exposed to the drug (since it determines the fraction of surviving susceptible genotypes). This is a black and white outcome, and it seems that there is little to understand beyond this obvious reasoning. Yet, in natural / real conditions (*i.e.* unlike in ecotoxicological tests performed in the laboratory), it is not very clear which dose of the drug is relevant. Concentrations of drugs vary at different temporal and geographical scales, within bodies (Levison and Levison 2009), within microhabitats, within regions *etc.* (Thiele-Bruhn 2003; Depledge 2011). Notwithstanding, it is often assumed that a resistance mutation is associated with a selection coefficient measuring the rate at which it is expected to change in frequency in populations, like any other beneficial mutation. This selective advantage is not easy to estimate in the field, but is often thought to represent an inherent property of the mutation itself. However, this advantage must depend in some ways on the exposure to the drug, which is an environmental variable, and not a property of the mutation.

### The context dependence of fitness effects

This point leads to a very simple and obvious idea: the selective effects of mutations depend on ecological conditions. This conclusion is somewhat trivial (Bell 2008), but it contradicts the naïve view that tends to essentialize the properties of mutations or genotypes (*i.e.* that characterize mutationnal properties as intrinsic). There is no such thing as a single selective effect of a mutation or a genotype. A mutation is not inherently beneficial, deleterious, or neutral. Rather, it depends on the ecological conditions, the genetic background, and on other alleles considered in comparison. Even the ‘dominance’ of mutations in diploids can be highly context and trait dependent (Bourguet *et al.* 1996; Manna *et al.* 2011). It is important to reiterate this point for several reasons. The first is that it tends to be simply ignored, even if this context dependence is never directly challenged. For instance, in molecular evolution, it is customary to assign a mutation, without further specifications, in three categories: deleterious, neutral or beneficial (Eyre-Walker and Keightley 2007). Secondly, it makes little sense to study the selective effects of mutations while neglecting the diversity of real ecological conditions and the diversity of genetic backgrounds. Of course, it is always possible to think in terms of average, and this is the usual view when dealing with the diversity of genetic backgrounds in sexual populations (where recombination ensures that mutations “experience” effectively the range of possible backgrounds, Lande 1983; Chevin and Hospital 2008). Yet, for ecological variation, taking such an average may be very misleading. For example, a resistance mutation can spread somewhere (in a treated environment), but not elsewhere (in a non-treated environment), generating a situation of polymorphism (Jain and Bradshaw 1966; Suckling and Khoo 1993; Carrière *et al.* 1994; Guillemaud *et al.* 1998; Lenormand *et al.* 1998, 1999; Lenormand and Raymond 2000; Neve and Powles 2005; Labbé *et al.* 2009). Differences of effects across environments matters, not only their average, provided that the spatial scale of dose variation is larger than the scale of dispersal (Lenormand 2002) or that temporal fluctuations occur at scales exceeding generation time (e.g. Cvijović *et al.* 2015).

### The cost of resistance

In presence of such heterogeneity with both treated and non-treated environments, the concept of ‘cost of resistance’ becomes important. This cost is defined as the selection coefficient of resistance mutations in absence of treatments (or similarly in absence of predator, parasite or pathogens when considering resistance in the context of biotic interactions). The idea of a “lower adaptive value” of resistant genotypes in the absence of treatment can be traced back quite far (e.g. in Dobzhansky 1951). The term “reproductive disadvantage” or no specific term is used in this context in the 60s and 70s, in empirical or theoretical papers (Abedi and Brown 1960; Gillespie 1975; Hickey and McNelly 1975; Antonovics 1977; Georghiou and Taylor 1977; Curtis *et al.* 1978). The term ‘cost of resistance’ is relatively recent in comparison and started to be widespread only in the 80s, especially in the context of plant resistance to herbivores (Windle and Franz 1979; Leonard and Czochor 1980; Simms and Rausher 1987) or bacteria resistance to phage (Lenski 1988). It is in particular used in the influential paper of Anderson and May (1982) on coevolution and resistance to pathogens. Curiously, when dealing with resistance evolution in an abiotic context (*i.e.* to pesticides or antibiotics), the term ‘cost of resistance’ still refers in the 80s to the economic, not the evolutionary, cost of resistance (*i.e.* the extra monetary cost due to increasing pesticide dosage consecutive to resistance evolution), even in papers by May, referring to evolutionary cost as ‘back selection’ (May and Dobson 1986). The first mention of an evolutionary cost of resistance in an abiotic context seem to be in Jacobs *et al.* (1988) about herbicide resistance, although the term “cost of tolerance” was used in studies of heavy metal tolerance in plants a bit earlier (Bradshaw 1984; Wilson 1988). The cost of resistance was then intensely investigated in the 90s (Bergelson and Purrington 1996; Andersson and Levin 1999 for reviews), with several methodological improvements (e.g. separating costs from effect of linked variation). By the end of the 90s, most pesticide resistance management models included the cost of resistance, *i.e.* the fact that resistance mutations could be selected against in non-treated environments (Lenormand and Raymond 1998). As this short historical overview shows, the concept of cost of resistance is relatively recent and was not used before the 80s. In particular, all the paper on local adaptation, clines and all the field of ecological genetics developed before the 80s without the need to refer to this concept. For instance, in classical models such as Levene model (Levene 1953) or cline models (Haldane 1948; Nagylaki 1975; Endler 1977), it was sufficient to talk about the selective effects of alleles in different environments. Why was the concept of cost of resistance introduced? At first sight, the concept seems asymmetrical and not very useful. For instance, nobody talks of the “cost of susceptibility” in treated environments, although it would be equally legitimate.

A likely reason is that the concept was helpful to bring attention to the fact that a mutation could be both beneficial or deleterious depending on circumstances, something well known in ecological genetics but somewhat ignored in resistance studies. It helped introduce some ecology in the understanding of the fitness effect of resistance mutations. This can have important consequences as the cost of resistance is a powerful force that can keep resistance in check (see e.g. Curtis *et al.* 1978 for an early model). Considering cost is also very important to predict the dynamics of resistance mutations in heterogeneous (treated and non-treated) environments.

A second reason is that the concept of cost was tightly associated to the notion of trade-off among traits, an idea borrowed from life history theory. When resistance is viewed as a trait, or a “defense” function, it is natural to consider that it may trade-off with other organismal traits and functions (e.g. in terms of resource). In other words, resistance mutations should influence many traits, *i.e.* have pleiotropic effects. Besides resistance, the variation of all these traits is likely to be deleterious, and therefore represent a ‘cost’, since these traits were previously optimized by natural selection. This connection with life history theory is entirely explicit in the first papers mentioning the concept (Simms and Rausher 1987; Lenski 1988; Wilson 1988; Smith *et al.* 1991; Bergelson and Purrington 1996) and led to idea that the cost of resistance was caused by the pleiotropic effects of resistance mutations. Although the idea of trade-off among traits was initially present in this interpretation, it is often forgotten today: resistance mutations are simply viewed as ‘pleiotropic’. Naturally, only considering pleiotropy, it seems very natural to think that the cost can evolve to be reduced, or even eliminated (which is also directly suggested by the word ‘cost’ itself). For instance, one could consider that compensatory evolution should attenuate these unwanted pleiotropic effects and restore optimal values for all traits, eliminating the cost. At first sight, the best proof for this reasoning is that cost-free resistance mutations are sometimes found along with the existence of modifier loci mitigating or even eliminating costs (McKenzie *et al.* 1982; Davies *et al.* 1996; Lenski 1998; Andersson and Levin 1999; Andersson and Hughes 2010; Melnyk *et al.* 2015).

Today, the term ‘cost of resistance’ is widely used, but the concept suffers from several ambiguity that can be understood in the light of this short history. First, the concept seems unnecessary to study adaptation in different environments (it would be sufficient to simply consider fitness effects in each environment). It also introduces an asymmetry, which is quite arbitrary, and somewhat misleading (susceptibility too is costly). Second, it conflates effects across traits (pleiotropy) and effects across environments, which can also be misleading. Third, the word ‘cost’, still reflects an essentialization of mutation/genotypes. The deeply engrained view “one mutation – one fitness effect” was not really challenged by the introduction of the ‘cost’ idea. It was merely replaced by the idea that one resistance mutation corresponded to two important characteristics: its benefit and its cost.

For these reasons, we think that the concept of cost of resistance presents important shortcomings, to the point, that it is now becoming a problem and hindrance to conceptually clarify the process of adaptation. We now try to explain these issues in more details.

### Costs of resistance are not equivalent to pleiotropic effects

The first problem with the concept of cost is its interpretation in terms of pleiotropic effects of mutations. To be very clear, a simple situation is sketched below where adaptation is represented by the optimization of many traits simultaneously, like in Fisher’s model of adaptation or multivariate models of stabilizing selection in quantitative genetics (Lande 1980; Hartl and Taubes 1998; Orr 1998; Martin and Lenormand 2006a), but with two environments (Martin and Lenormand 2006b, 2015). Figure 1 uses only two traits, which is sufficient to make the argument and discuss the issue of pleiotropy. It is straightforward to relate this type of fitness landscape model to the more traditional one-dimensional ‘dose-response’ models (see Box 1). Representing evolution of resistance as convergence to a phenotypic optimum has received some empirical support (Bataillon *et al.* 2011; Sousa *et al.* 2012; Harmand *et al.* 2017, 2018) and may capture well the dynamics of adaptation.

**Figure 1.**
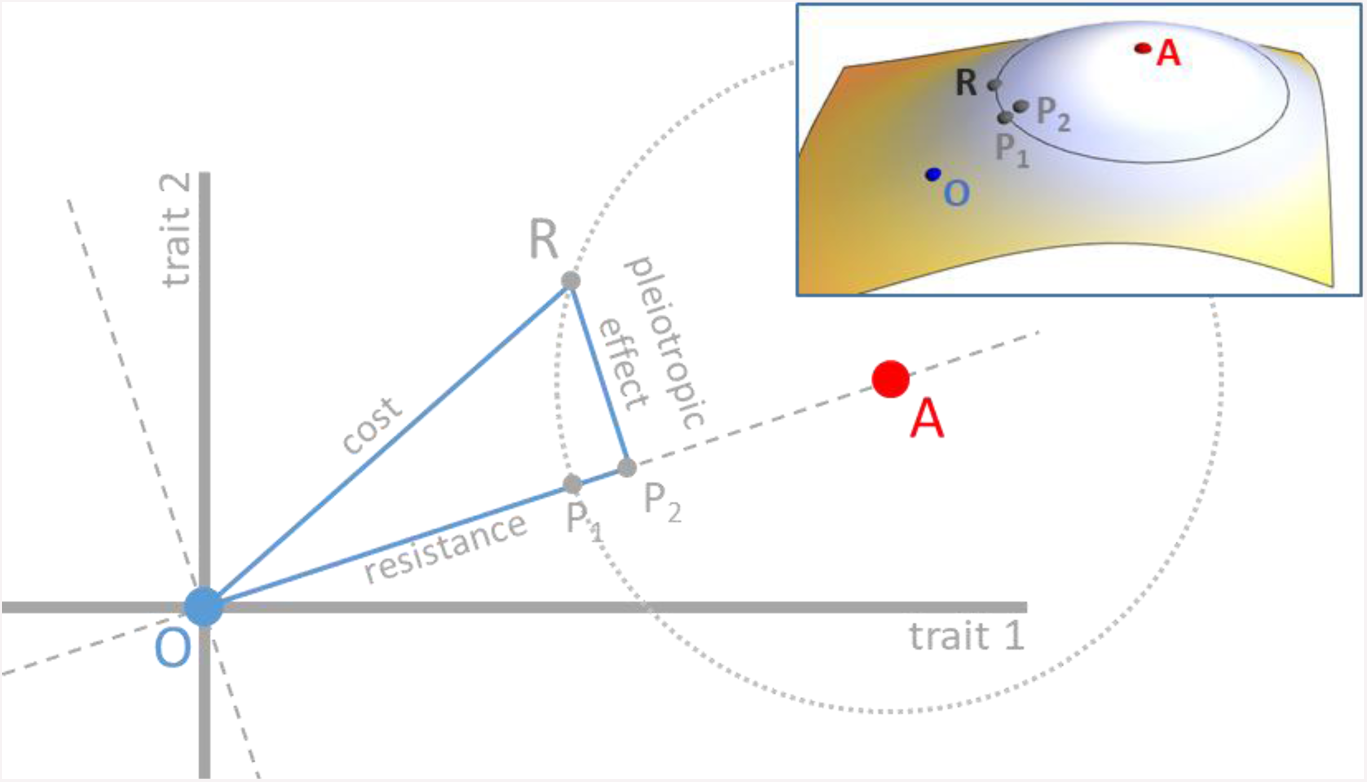
Graph of treated and non-treated environments, with distinct phenotypic requirements (phenotypic optima A and O, respectively) in a two-trait space. Assuming a wild-type positioned in O, the resistance mutation R brings the phenotype closer to A. Relative to the wild type, R is therefore a beneficial mutation in the treated environment. Its cost is usually defined as its fitness in the non-treated environment relative to the wild type, which depends on the distance between R and O on the figure. Note that the cost (and all fitness measures on this figure, and similar figure below) depends on Euclidian distances in phenotypic space, and a mapping function converting this distance to fitness (*i.e.* the cost is not distance OR, but a monotonic function of this distance). The mapping is left implicit on the figure, but can be thought as a third orthogonal axis representing fitness for each trait 1 – trait 2 combination, which defines a “fitness landscape”. The coloured inset figure represents such a fitness mapping in 3D. Fitness values, when projected on the phenotypic space correspond to isofitness curves (like altitude on a geographic map is indicated by contour lines). For instance all phenotypes on the light grey circles have the same fitness than mutation R in the treated environment (optimum A). The direction of the two optima (OA axis) defines a phenotypic trait of ‘resistance’. Variation of trait(s) orthogonal to this axis may be defined as pleiotropic effects. Point P_1_on OA axis is such that AR = AP_1_. It represents the phenotypic point that would confer the same fitness in the treated environment compared to the mutation R, but that would only alter the phenotype in the exact direction of the optimum A. Point P_2_is the orthogonal projection of R on OA axis. It represents the phenotypic point that would be reached or if all the pleiotropic effects of the mutation R were compensated (e.g. by subsequent compensatory mutations).

Let us label “O” the optimal phenotype in the non-treated environment and “A” the phenotypic optimum in the treated environment. For simplicity, we can assume that fitness monotonously declines with the (Euclidian) distance from the peak in any given environment. Again, this simplified model could be more specific (with a particular mapping of distance to fitness) or complex but the core argument does not require the use of more complex situation or assumptions. We can also assume that the wild type is very close to O, as one would expect from the effect of past selection in absence of drug, and represents a resistance mutation by a vector pointing from O to R, where R is a phenotype closer to A than to O. The mutant is beneficial (relative to the wild type) in the treated environment because the distance AR is smaller than the distance AO. The difference between these two distances scales with the selection coefficient of the resistance mutation in the treated environment. The OA axis, by definition, represents the phenotypic direction of the ‘resistance’ phenotype. Point P_1_ on this axis such that AR = AP_1_allows representation of the phenotypic point that would confer the same benefit in the treated environment compared to our resistance mutation R, but that would only alter the phenotype in the exact direction of the optimum A. In other words, the distance OP_1_scales with the selective advantage of the resistance mutation in the treated environment, relative to the wild type. The cost of resistance depends on the distance OR, as it is defined as the fitness effect of the resistance mutation in the non-treated environment (again, relative to the wild type). What about the pleiotropic effects? Here, the OA axis represents the phenotypic axis of resistance and therefore, the orthogonal direction represents all other traits (here, there is only one other trait, because we consider only a two dimensional trait space but in with *n* phenotypic traits, there would be *n* − 1 such traits). Hence, the ‘other’ pleiotropic effects all project on these axes orthogonal to OA. Furthermore, note the point P_2_, the projection of point R on the OA axis. The vector RP_2_ represents the pleiotropic effects of the resistance mutation. Should these effects be totally compensated, the phenotype would be in P_2_ and it would indeed enjoy a greater fitness in both the treated and non-treated environments (since AP_2_< AR=AP_1_and OP_2_ < OR, respectively).

This simple geometric argument indicates several things. First, pleiotropic effects and the ‘cost of resistance’ are two different things – biologically and geometrically – contrary to what is usually considered. Pleiotropic effects will be eventually compensated through the well-known process of “amelioration” when the population reaches the phenotypic optimum after remaining exposed to the treated environment and involves new resistance mutations, compensatory mutations or a mixture of mutations with the two properties (Cohan *et al.* 1994; Schrag *et al.* 1997; Lenski 1998; Levin *et al.* 2000; Schoustra *et al.* 2006; MacLean and Vogwill 2015). Cost evolution will be quite different, and may occur if the population is exposed, at least part time, to the non-treated environment (by evolution of plasticity or inducible response, e.g. Nguyen *et al.* 1989; Foucault *et al.* 2009). Second, the full compensation of these pleiotropic effects does not reduce the cost of resistance to zero. Indeed, biologically speaking, it is likely that acquiring ‘resistance’ requires changing at least one trait, and thus, this trait becomes suboptimal in the original environment for the resistant mutant. This is irreducible and corresponds to the idea that different phenotypic requirements necessarily involve the occurrence of a fitness trade-off. In any case, there is no reason to believe that the deleterious pleiotropic effects of a resistance mutation at a given drug dose are equal to the cost of resistance at dose zero. Finally, it points out that the fitness effects of mutations are not a fixed property of that mutation. It also depends largely on the environments (here the positions of optima). We discuss now this idea in more details.

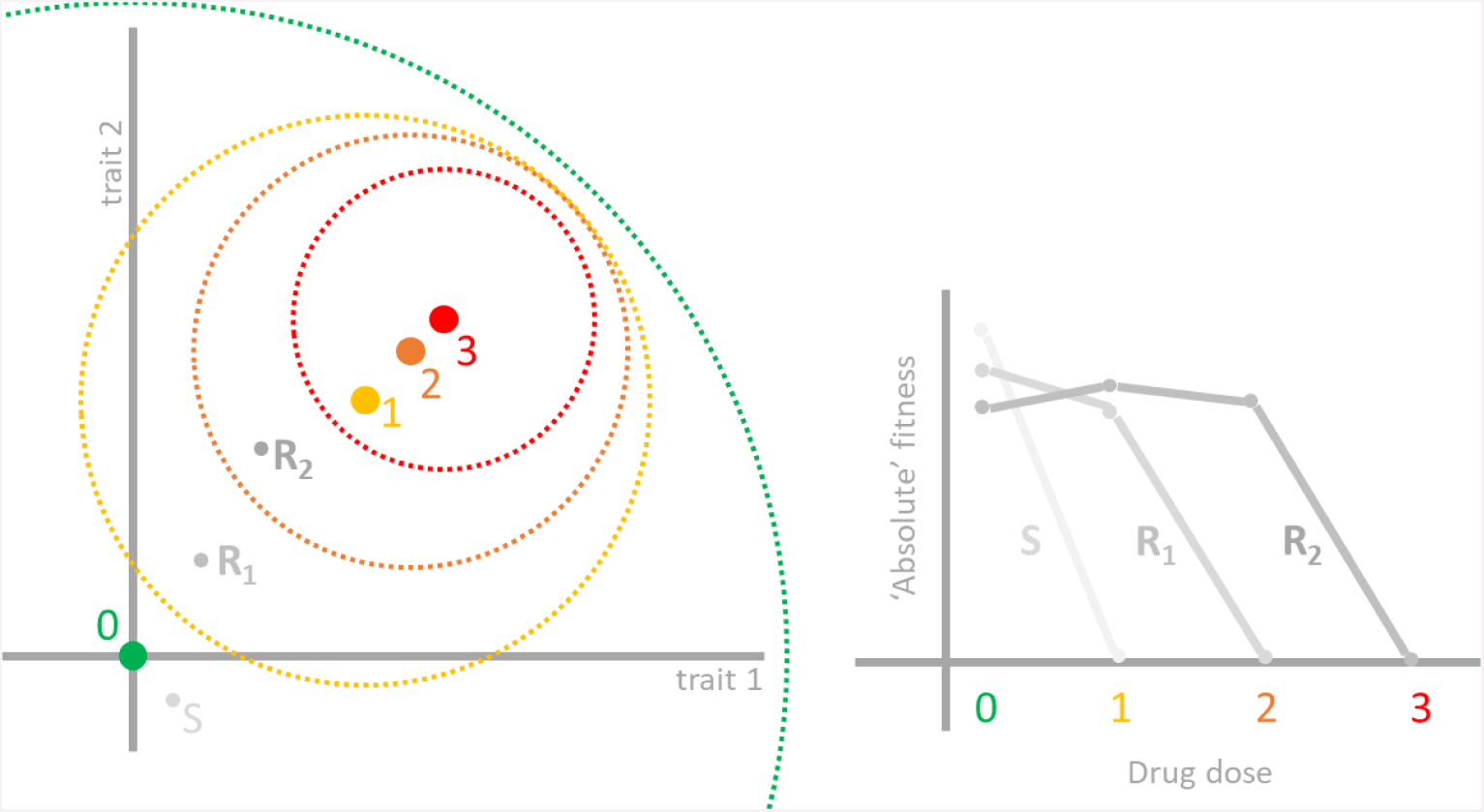

Box 1. Correspondence between a fitness landscape model and a drug-response model. On the left panel, a fitness landscape model is illustrated as in Figure 1 to 4, in a two-traits phenotypic space. Four environments are represented with increasing concentration of a drug (with optima 0, 1, 2, 3). Environment with optimum 0 (in green) represents the environment without drug, while environment 1, 2, 3 represent increasing concentrations of the drug. In this model, fitness depends on the Euclidian distance to the optimum, and a mapping function (see inset of Fig. 1). It is possible to set a threshold value for absolute fitness below which a phenotype cannot persist / grow. This threshold is indicated by circles (the colour of the circle corresponds to the different environments). In many cases, resistant mutants can have a positive growth rate in absence of drug (while the reverse is not true: susceptible phenotype do not grow in presence of the drug). Hence the threshold contours will be often nested (but it is possible to imagine cases where this is not the case). In this representation, it is easy to see that an absolute fitness criterion (= being within the threshold contour) is not synonymous with adaptation (= being close to the optimum). The dot S represents the position of a susceptible phenotype, while R_1_ and R_2_ represent two resistance mutants. The right panel illustrates the ‘dose response’ curves relating, dose to absolute fitness, for each phenotype (S, R_1_, R_2_). For instance, R_1_ is within the threshold contour of dose 1, but not of dose 2 and 3. Hence, its dose-response is zero for doses above dose 1. Note also that R_1_ is further apart from optimum 0, compared to S. Hence, its absolute fitness is lower than that of S at dose 0. This correspondence shows that it is entirely possible (and straightforward) to relate fitness modes to more traditional one-dimensional dose-response models. Furthermore, since LD_50_, IC_50_, MIC or other ecotoxicological measures can be defined using dose response, they can also be defined in the fitness landscape model. Note however that these measures have been rightly criticized as being only partial fitness summaries (Regoes et al. 2004; Sampah et al. 2011; Wen et al. 2016). They are also often obtained in absence of competition, or often concern only a particular life stage. Note also that absolute measures of fitness are often more appropriate (than relative fitness) when dealing with the demography of the treated species (Day *et al.* 2015). However, relative fitness is in general more relevant to study environment specialization, where the “cost of resistance” matters.

### Resistance mutations do not have *“a cost”*

From an experimentalist perspective, defining a non-treated and treated environment is straightforward. You first select a given environment, then you can either add the drug or not. With this definition, it is possible to make a very clear and clean experiment demonstrating the effect of the drug, with a control. Yet, the problem is that there is virtually an infinite set of possible environments to start with. Which pair of treated/non-treated environments is relevant? This is difficult to know. It is difficult to represent the complexity of natural conditions in controlled experiment in the laboratory. Even trying to determine which environment corresponds to the environment to which an organism has been adapting to is challenging. For instance, using ‘absolute’ demographic performance to answer this question may not be reliable. For instance, habitat quality varies and can even obscure the relationship between ‘absolute’ measures of fitness and environment variables (Gallet *et al.* 2014). For instance, *E. coli* in the gut of humans evolved for a long time at 37°C, yet it grows faster at slightly higher temperatures in laboratory conditions (Gonthier *et al.* 2001). Hence, the variation in growth rate (often taken as an absolute measure of fitness) may not be used so easily to infer the environment where adaptation took place.

In fact, it is quite straightforward to see that the cost of a mutation will be different in varying non-treated environments. There is not “a” cost, but as many costs as there are different (non-treated) environments (Gassmann *et al.* 2009; Vila-Aiub *et al.* 2009; Angst and Hall 2013; Gifford *et al.* 2016), which may also be revealed by different compensatory evolution in different environments (Björkman *et al.* 2000). Worse, this cost of resistance may not even actually be positive, challenging the usage of the word “cost” itself (in the usual economic sense, a cost is necessarily positive, or it would not be a “cost” in the first place). In other words, the resistance mutation may be favourable in both the treated and non-treated environment, relative to the wild type (see examples in Kassen and Bataillon 2006; Melnyk *et al.* 2015). This can occur for many reasons, but globally will occur often when the wild-type is not well adapted neither to the treated nor to the non-treated environment (Martin and Lenormand 2015). Such a situation is illustrated on Figure 2. Let keep the position of the wild type and resistance mutant in O and R, respectively. Let’s also keep the position of the optimal phenotype in the treated environment (in A). Now, lets consider that the optimal phenotype in the non-treated environment is not O, as in Figure 1, but is B. Reporting the point P_3_ such that BP_3_= BR, we see that the distance to the non-treated optimum is greater for the wild type than for the mutant (*i.e.* BR < BO). The difference between BR and BO actually corresponds to OP_3_. When the resistance mutant is favourable in both the treated and non-treated environments (like in this example), the term ‘cost’ becomes confusing since it implies talking about a ‘negative cost’. It would also, in this case, be unclear to interpret costs as deleterious pleiotropic effects (since there is no deleterious effect in the first place). If costs are not pleiotropic effects and not even costly (meaning deleterious), the terminology and its usual interpretation start obscuring things instead of clarifying them.

**Figure 2.**
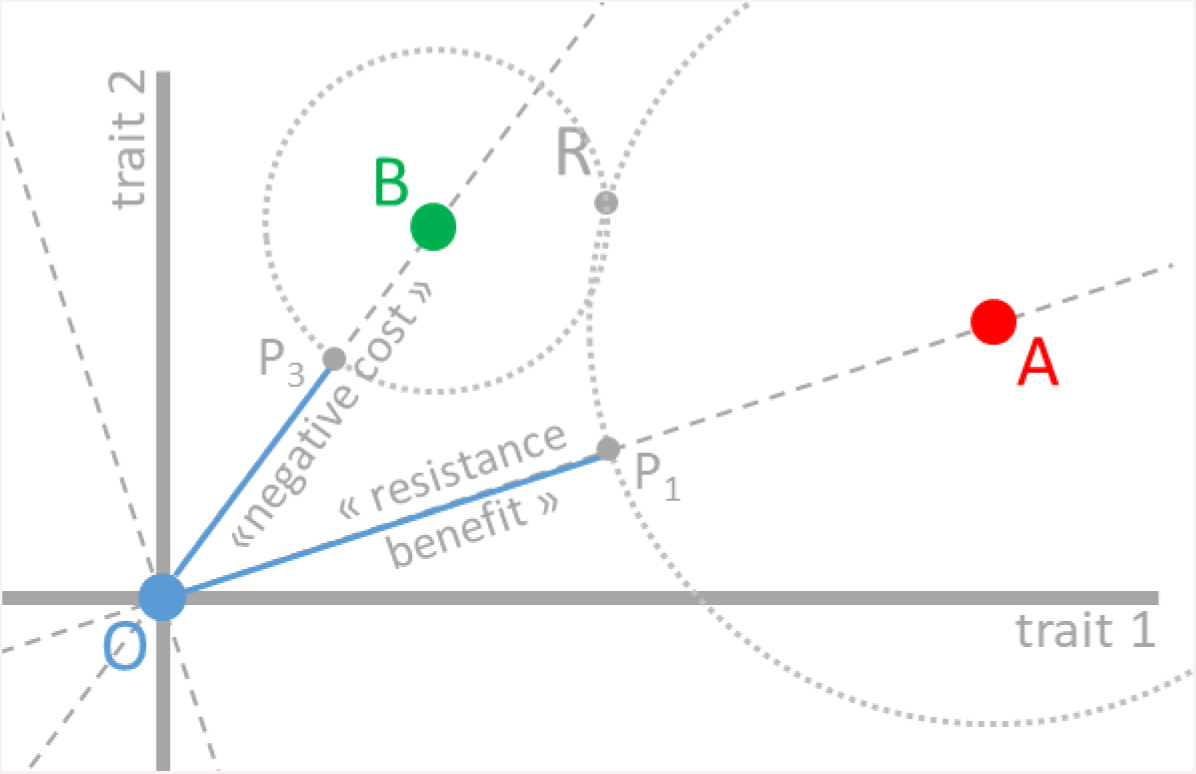
Graph of treated and non-treated environments, with distinct phenotypic requirements (phenotypic optima A and B, respectively) in a two-trait space. Relative to a wild-type positioned in O, the resistance mutation R brings the phenotype closer to A. As in Figure 1, R is therefore a beneficial mutation relative to the wild type O in the treated environment (with optimum A). However, its cost is now “negative” in the non-treated environment, as R is also closer to B compared to the wild type O. The point P3 is such that BP_3_ = BR. The distance to the non-treated optimum B is greater for the wild type O than for the mutant R (*i.e.* BR < BO). The distance difference between BR and BO corresponds to OP_3_. As in Figure 1, all fitness measures depend on the phenotypic distances illustrated and a mapping that could be represented as a third orthogonal axis representing fitness. This fitness axis is not shown. All phenotypes on the light grey circles have the same fitness than mutation R in the treated environment (optimum A, large circle) and non-treated environment (optimum B, small circle).

Such situation happens when RB ≤ OB *i.e.* when the wild-type and the resistance mutant are at least equally distant from the non-treated optimal phenotype. Because such situations are quite common (either because we cannot properly reproduce natural conditions experimentally, or because wild type genotypes are not well adapted to their environment), it is perhaps not very surprising that sometimes “cost free” resistance mutation or even “negative cost” are found. All these situations can occur but are unrelated to the pleiotropic effects of the resistance mutation (as defined in the previous section). Importantly, finding an absence of cost or even ‘negative costs’ do not indicate necessarily that there is no phenotypic trade-off between the treated and non-treated environment (and indeed, on the Figure 2, A and B are distinct points, indicating that there is a phenotypic trade-off, despite that no cost is detected). It can just indicate that the wild-type reference is not adapted well to the non-treated environment. At this point, one might argue that all this confusion arises because the mutation R was not a “resistance” mutation to begin with. The mutation R is beneficial in both the treated and non-treated environment, and so, it may be better interpreted as a beneficial mutation to the non-treated environment than as a ‘resistance’ mutation. For instance, if we are talking of a bacteria mutant showing this property in a laboratory test, we may want to say that the mutation R corresponds to adaptation to the “laboratory condition”, not really to the drug *per se*. But, then, how is a resistance mutation truly defined?

### What is a resistance mutation?

If resistance mutations cannot be defined by the fact that they are beneficial in the treated environment, relative to wild type (as we did up to now), then, how do we classify them? In fact, it might be possible to define them more specifically by saying that they are beneficial in the treated environment relatively to wild type, provided that the wild type is perfectly well adapted to the corresponding non-treated environment. In principle, this definition makes sense, as it avoids conflating adaptation to conditions that are shared by both treated and non-treated environments, with adaptation to the drug itself. Yet, with this definition, the mutation R illustrated on Figure 2 would still be a beneficial mutation in the treated environment. R is closer to A (the optimum with the drug) than would a wild type well adapted to the non-treated environment (distance RA < AB). In such a case, the phenotypic direction corresponding to resistance would be the AB axis (and the pleiotropic effects best defined on axes orthogonal to AB). This would be in general clearer and more insightful. With such a definition, it is possible to distinguish mutation R and R’ on Figure 3 for instance. Both would be beneficial in both treated and non-treated environment, relative to a wild-type in O, but only R would be beneficial in the treated environment relative to a wild-type in B.

**Figure 3.**
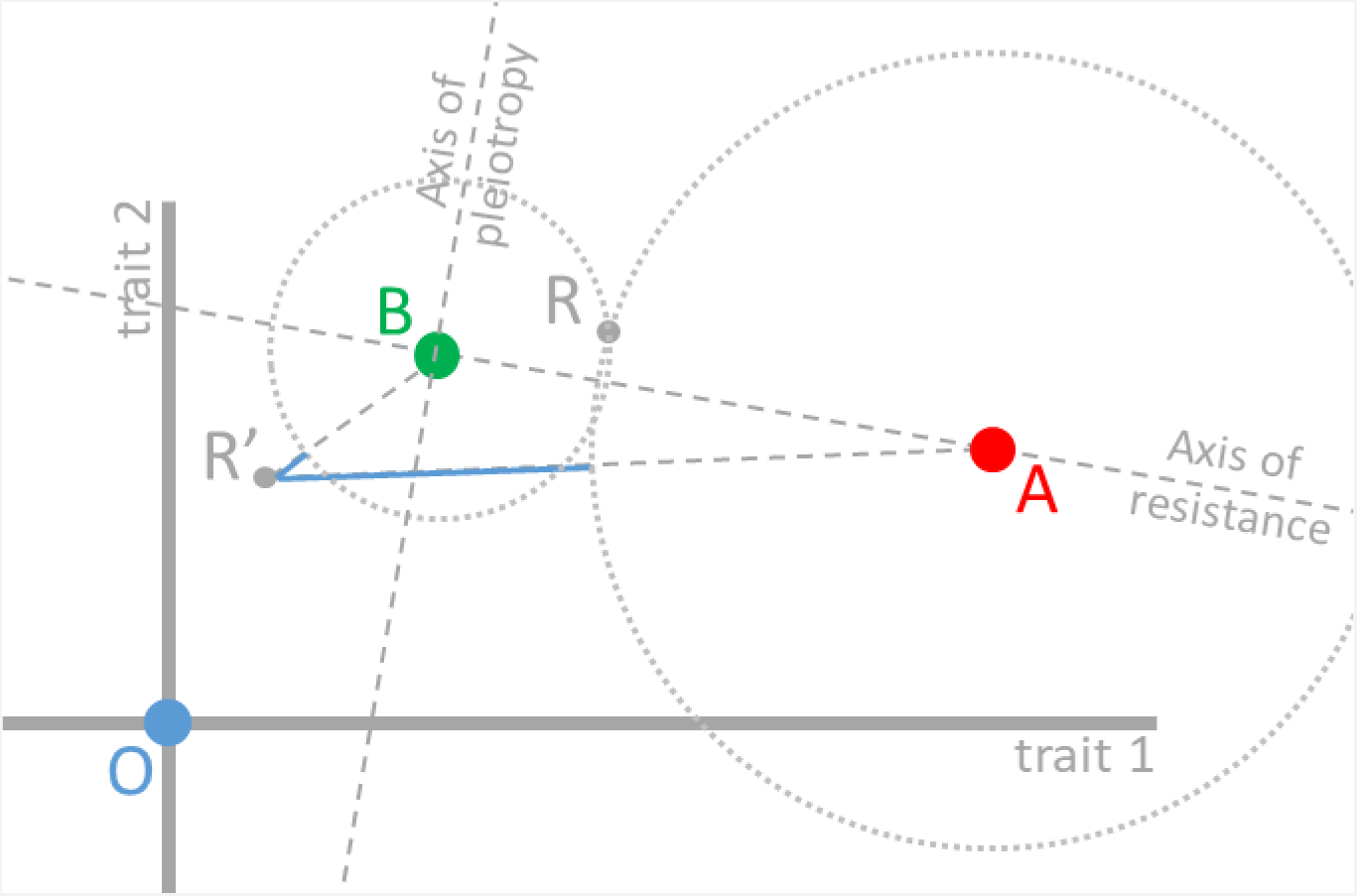
Sketch of treated and non-treated environments, with distinct phenotypic requirements (phenotypic optima A and B, respectively) in a two-trait space. Relative to a wild-type positioned in O, both the resistance mutation R and R’ brings the phenotype closer to A. As in Figure 1, they are therefore both beneficial mutations, to the treated environment, relative to the wild type O. However, if resistance is defined by comparing to a wild-type well adapted to the non-treated environment (*i.e.* to a wild type located on B), then only mutation R would be beneficial, and therefore could qualify as being a resistance mutation. With this comparison, costs would also be better defined (*i.e.* would always be positive). As with Figure 1 and 2, fitness depends on phenotypic distance and a mapping, which is not illustrated but would correspond to a third orthogonal axis.

The problem is that measures of fitness will be made against a wild-type and it may not be straightforward to determine whether this wild type is well adapted to the non-treated environment (but not impossible, as e.g. in cases of experimental evolution in the laboratory where the wild type can be chosen as the type that evolved in the non treated environment for a long time). Without this knowledge, it may be very difficult to classify mutations that are specifically beneficial in the treated environment (*i.e.* “resistance” mutations as defined here), versus mutations that are beneficial in both treated and non-treated environments (see e.g. Marcusson *et al.* 2009 for an example of this problem).

There is yet another difficulty lurking in the vast range of possible natural situations. Just as there are many non-treated environments, there may be many treated environments as well. In particular, any drug may be added in different quantities or concentrations to any given environment. Here resides a problem, which is rarely addressed in studies on resistance: different drug concentrations can correspond to different intensity of selection (Milesi *et al.* 2016), but they can also correspond to different optimum phenotypes (Harmand *et al.* 2017). This possibility needs to be demonstrated empirically (as in Harmand *et al.* 2018), and cannot be ignored *a priori*. Admitting that natural processes have some degree of continuity, it is likely that differing doses end up corresponding to different phenotypic optima. Indeed, it is difficult to conceive that adding a vanishingly small quantity of drug suddenly shifts away phenotypic requirements, and that further increases in doses only change the selection intensity around that shifted phenotypic peak.

If different drug doses correspond to different optima (say A_1_and A_2_on Figure 4), it is fairly easy to imagine two mutations R_1_and R_2_that would qualify as resistance mutations, in each of the two environments, but not in the other. On Figure 4, R_1_is a resistance mutation for drug dose 1 (optimum A_1_), but not for drug dose 2 (optimum A_2_), and reciprocally for R_2_. This difference does not arise because these mutations have different “costs”. In fact, the two mutations illustrated on Figure 4 have the same cost (the same fitness in non-treated environment B). This situation occurs because phenotypic requirement is different with different doses, such that a phenotypic change can be favourable in one environment (here one dose), but not another. ‘Resistance mutations’ can differ not only in their benefit and costs at a given drug dose, but also in their fitness effects at other doses. When optima for doses are different, there are many underlying trade-offs, which are not necessarily caused by differing costs in the non-treated environment (Harmand *et al.* 2018). Trade-offs among doses are not captured by studying costs. A common view is that mutations conferring strong resistance (*i.e.* resistance to a high dose) may carry strong costs (Melnyk *et al.* 2015), explaining perhaps why they may not be beneficial at a lower dose. This may well be true, but not necessarily (Harmand *et al.* 2017). A mutation favourable at high dose may be deleterious at low dose, irrespectively of its cost. This is the case illustrated on Figure 4. Studying “cost” and “benefit” at one particular dose may give the illusion that all trade-offs are understood. In fact, this is not the case: all trade-offs among doses will be missed.

**Figure 4.**
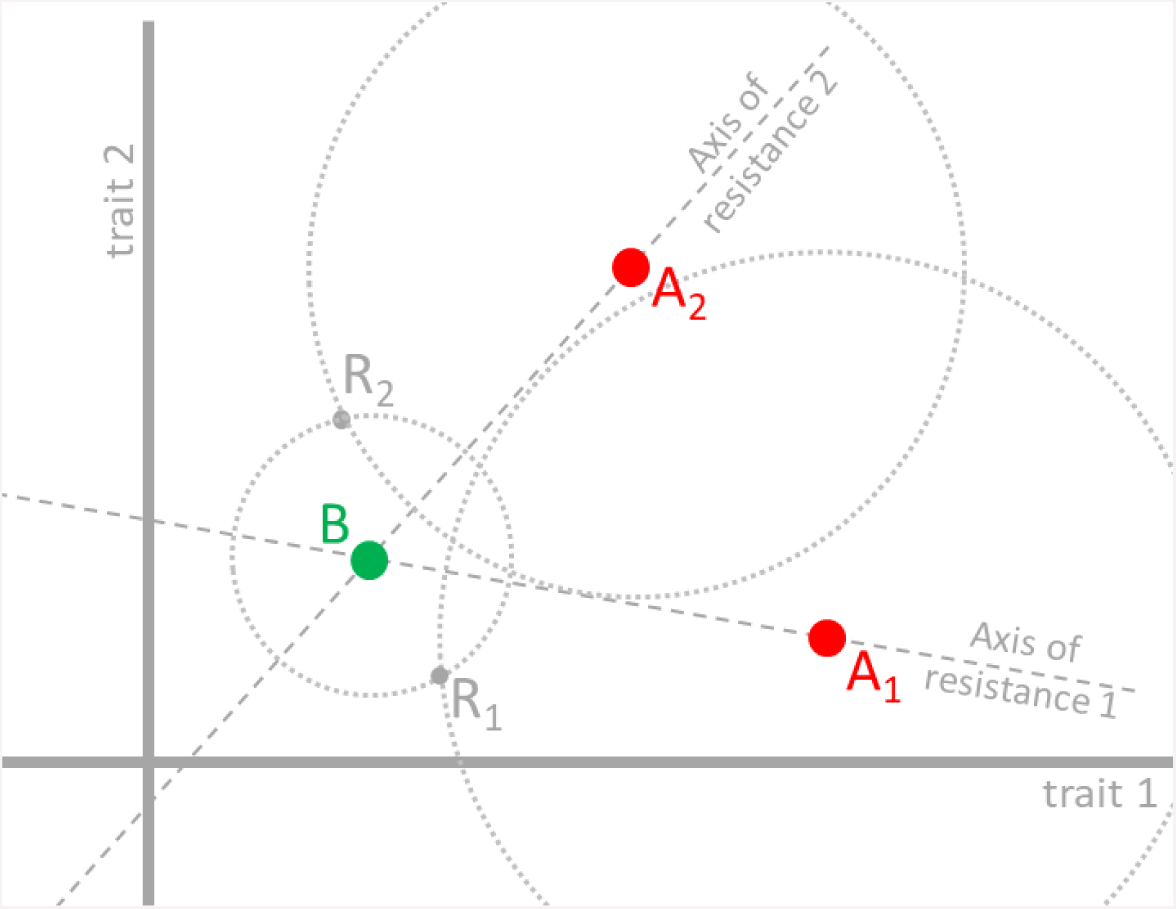
Sketch of a non-treated environment (phenotypic optimum in B) and two treated environments, with different doses of drug (optima A_1_ and A_2_). Two mutations are illustrated, with fitness effects compared to a wild type well adapted to the non-treated environment (*i.e.* located at B). Using the definition from Figure 3, R_1_ is a resistance mutation with respect to environment A_1_, but not with respect to environment A_2_, and reciprocally for mutation R_2_. Nevertheless, both mutations have the same cost. As with Figure 1 to 3, fitness depends on phenotypic distance and a mapping, which is not illustrated but would correspond to a third orthogonal axis.

### Summary, conclusion and practical implications

Taking into account the ‘cost of resistance’ has been a major progress because it is essential to distinguish the fitness effects of resistance mutations in treated versus non-treated environments. However, this cost is also very expensive conceptually as it is associated with too many simplifications, to the point that it may even be misleading. In practical terms, should the term ‘cost of resistance’ be avoided? The term is extremely widespread and in many cases, it has at least the merit to attract the attention to the fact that fitness effects are different in different environments. In most cases and depending on context, the more robust concepts of fitness trade-off across environments or pleiotropy could be used, although the concision of the expression “cost of resistance” will be difficult to match. We hope that this perspective will help correcting some of the sloppy usage of the term and dismiss the implicit expectations based on this terminology.

First, the interpretation of “cost” in terms of pleiotropic effects is very unclear. Pleiotropic effects may be better defined as effects projected on phenotypic axes orthogonal to resistance phenotype, than in terms of fitness effect in the non-treated environment (see Figure 1). In practical terms, this indicates that the fitness effects across environments should be better distinguished from pleiotropic effects. Pleiotropy is not necessarily the cause of fitness trade-off. A single trait can present different optimal values in different environments and primary resistance traits can exhibit a trade-off without having a pleiotropic effect. This clarifies our expectation about the process of adaptation: compensatory evolution is not expected to systematically reduce the “cost of resistance” to zero.

Second, there are as many costs as there are non-treated environments. The idea that there is one cost associated with a resistance mutation is an extreme and naïve simplification. In practice, this indicates that studying the fitness effects of resistance mutations “in the wild” (*i.e.* beyond the simplified laboratory conditions) is very important.

Third, costs of resistance are ill defined when several precautions are not taken. For instance, failing to measure costs relative to a well-adapted wild type to the non-treated environment can lead to absurd notions, such as ‘negative costs’ that serve no conceptual clarification (Figure 2). In practice, these precautions can be very difficult to meet, as the degree of adaptation of the wild type to the non-treated environment will be in general difficult to assess, but steps can be taken in that direction (e.g. by carefully choosing the reference genotype and the reference environment). In any case, this issue is important to keep in mind when interpreting results.

Fourth, the concept of cost leads to an oversimplified and often erroneous view of trade-offs across environments. Finding costs equal to zero (*i*) cannot be used to say that there is no trade-offs between treated and non-treated environments; (*ii*) cannot be interpreted by saying that pleiotropic effects were compensated; (*iii*) completely misses the possibility that fitness trade-off may occur among different doses. In practice, finding a mutation with no “cost” (or a “negative cost”), should not be surprising. It does not prove the existence of “Darwinian demons”; it should not be used to imply that there are no trade-off across environments and therefore that resistance management strategies will necessary fail. As we have shown, this finding can simply result from a particular choice for the reference genotype and environment. Considering fitness effects across the range of possible doses is also important beyond the simplified conditions of most lab-based ecotoxicological tests and ecotoxicological fitness proxies such as LD50, MIC, dose responses *etc*. Importantly, finding that resistance mutations are not favourable at all doses cannot be attributed to their “cost”. For instance, high resistance mutations may not be beneficial at low doses, not because they have a high cost, but simply because they do not match phenotypic requirements that are optimal at low dose. Here again, the concept of cost leads in practice to biased expectations. Finally, observing that a resistance mutation does not decrease in frequency after an arrest of treatment, is not proving that there is “no cost” or no trade-off across environments. Other causes of frequency changes must be first investigated (*i.e.* effect of gene flow, effect of residual or hidden treatments, drift, indirect selection) as well as possible ascertainment biases (low power to detect slow frequency change).

Overall, it may be safer in most cases to simply discuss and measure the fitness effects of mutation in different environments and to carefully consider the role of the reference genotype/phenotype when interpreting relative fitnesses. All these points hold for many other situations of adaptation besides resistance, but this is perhaps where the vocabulary and the conceptual issues are the most acute and widespread. Differences among selective conditions and the occurrence of pleiotropy are both important ideas in evolutionary ecology. However, they cannot be solely summarized by assigning resistance mutations a ‘benefit’ and ‘a cost’ and essentialize their properties. There is a problem with a reduction of evolutionary thinking to a cost-benefit thinking, with “fitness” as a universal currency, valid regardless of ecological conditions. Although fitness is a general concept and a universal currency for adaptation, different conditions entail different fitnesses and possibly different phenotypic requirements (different adaptations). The vocabulary that we use should not oversimplify these ideas.

## Acknowledgments

We thank Helen Alexander, Danna Gifford, Inês Fragata and Claudia Bank for comments on the manuscript. We thank G. Martin for insightful discussions.

